# Expert Specification of the ACMG/AMP Variant Interpretation Guidelines for Genetic Hearing Loss

**DOI:** 10.1101/313734

**Authors:** Andrea M. Oza, Marina T. DiStefano, Sarah E. Hemphill, Brandon J. Cushman, Andrew R. Grant, Rebecca K. Siegert, Jun Shen, Alex Chapin, Nicole J. Boczek, Lisa A. Schimmenti, Jaclyn B. Murry, Linda Hasadsri, Kiyomitsu Nara, Margaret Kenna, Kevin T. Booth, Hela Azaiez, Andrew Griffith, Karen B. Avraham, Hannie Kremer, Heidi L. Rehm, Sami S. Amr, Ahmad N. Abou Tayoun, ClinGen Hearing Loss Clinical Domain Working Group

## Abstract

Due to the high genetic heterogeneity of hearing loss, current clinical testing includes sequencing large numbers of genes, which often yields a significant number of novel variants. Therefore, the standardization of variant interpretation is crucial to provide consistent and accurate diagnoses. The Hearing Loss Variant Curation Expert Panel was created within the Clinical Genome Resource to provide expert guidance for standardized genomic interpretation in the context of hearing loss. As one of its major tasks, our Expert Panel has adapted the American College of Medical Genetics and Genomics/Association for Molecular Pathology (ACMG/AMP) standards and guidelines for the interpretation of sequence variants in hearing loss genes. Here, we provide a comprehensive illustration of the newly specified ACMG/AMP hearing loss rules. Three rules remained unchanged, four rules were removed, and the remaining twenty-one rules were specified. Of the specified rules, four had general recommendations, seven were gene/disease considerations, seven had strength-level specifications, and three rules had both gene/disease and strength-level specifications. These rules were further validated and refined using a pilot set of 51 variants assessed by curators. These hearing loss-specific ACMG/AMP rules will help standardize variant interpretation, ultimately leading to better care for individuals with hearing loss.

**GRANT NUMBERS:** Research reported in this publication was supported by the National Human Genome Research Institute (NHGRI) under award number U41HG006834.

## INTRODUCTION

Sensorineural hearing loss (SNHL) is the most common congenital sensory deficit, with an estimated prevalence of 2-3 cases per 1000 individuals (Morton & Nance, 2006). Approximately half of SNHL in children is due to genetic causes, with 70% being nonsyndromic and 30% syndromic. Although variation in the *GJB2* gene is the most common cause of nonsyndromic genetic SNHL in many populations, there is high genetic and allelic heterogeneity, with over 100 implicated genes (Hereditary Hearing Loss Homepage http://hereditaryhearingloss.org) (Abou Tayoun et al., 2016; Alford et al., 2014).

Given the substantial genetic contribution to SNHL, a clinical genetics evaluation, including comprehensive sequencing, is recommended as part of a standard of care diagnostic work-up. Results of genetic testing can inform clinical management, particularly if a genetic syndrome is identified before the onset of additional clinical manifestations (Alford et al., 2014). The interpretation of sequence variants is a critical component of an accurate genetic diagnosis, and discrepancies in variant interpretation and classification have been well-documented and can have serious implications for patient care (Amendola et al., 2016; Booth et al., 2015; Booth, Kahrizi et al., 2018; Harrison et al., 2017). The large number of genes sequenced during hearing loss genetic testing routinely yield a large number of novel variants, exacerbating the interpretation challenge. Illustrating the extent of discrepancies in variant interpretation, as of 04/12/2018, 257 (8.1%) of the 3162 sequence variants in the 9 major hearing loss genes represented in our variant pilot (*USH2A*, *GJB2*, *SLC26A4*, *MYO7A*, *KCNQ4*, *TECTA*, *MYO6*, *COCH*, and *CDH23*) had conflicting interpretations in ClinVar.

In an effort to foster accurate interpretation and to reduce discrepancies across laboratories, the American College of Medical Genetics and Genomics (ACMG) and the Association for Molecular Pathology (AMP) published recommendations and guidelines for the interpretation of sequence variants (Richards et al., 2015). The ACMG/AMP guidelines contain several types of evidence that are weighted and categorized. Although these guidelines were intended to be used universally for all Mendelian disorders, certain criteria require gene- or disease-specific knowledge, a lack of which has been shown to contribute to discrepancies in variant interpretation (Pepin et al., 2016). In response to these issues, the NIH-funded Clinical Genome Resource (ClinGen, http://www.clinicalgenome.org) was launched as a centralized resource to provide guidance and tools for defining the clinical validity of gene and variant contributions to disease. Specifically, the ClinGen Hearing Loss Clinical Domain Working Group (HLWG) was established in 2016 to form Expert Panels to evaluate gene-disease associations, as well as standardize variant interpretation in hereditary hearing loss and related syndromes.

Here, we document the work of the Hearing Loss Variant Curation Expert Panel, hereafter referred to as the HL-EP. We adapted the ACMG/AMP guidelines for variant interpretation in the context of hearing loss. These specified rules were developed using the most common genes and syndromes that contribute to genetic hearing loss (HL); with the intent that most specifications could be used broadly across the many associated genes. These hearing loss-specific guidelines will be used in the future by the HL-EP to submit variant interpretations to ClinVar as an “expert panel” submitter, with the goal of resolving discrepancies, moving VUSs towards Benign or Pathogenic and increasing the confidence in HL variant classifications in ClinVar.

## MATERIALS AND METHODS

### Hearing Loss Clinical Domain Working Groups

As a ClinGen Clinical Domain Working Group (CDWG), the Hearing Loss CDWG aims to create a comprehensive, standardized knowledge base of genes and variants relevant to syndromic and nonsyndromic HL. Members were identified and recruited based on their expertise in hearing loss, and are representative of diverse institutions worldwide, spanning Asia, Australia, Europe, and North America. Members include otolaryngologists, clinical geneticists, molecular geneticists, ClinGen biocurators, clinical researchers, and genetic counselors from over 15 institutions. The Hearing Loss CDWG has so far formed two major efforts defined by ClinGen as a Gene Curation Expert Panel, and a Variant Curation Expert Panel (HL-EP).

### Specifications of the ACMG/AMP Guidelines

A smaller task team utilized biweekly conference calls in addition to email correspondence to review, specify, and reach consensus for each of the rules within the ACMG/AMP guidelines. Typically, 15-20 members from the Laboratory for Molecular Medicine (LMM), Children’s Hospital of Philadelphia (CHOP), ARUP Laboratories, Molecular Otolaryngology and Renal Research Laboratories (MORL), and Mayo Clinic participated on each call. Members with specific areas of expertise (e.g. Pendred syndrome) were also contacted for relevant topics.

The chairs, ClinGen biocurators, and coordinator of the HL-EP also utilized weekly conference calls for additional planning and preparation. For the larger CDWG, a quarterly conference call was used to present specifications finalized by the smaller group and to obtain final approval from all members. Upon specifying the ACMG/AMP rules, conference calls and email correspondence were used for the variant pilot using a set of 51 variants from nine common HL genes that represent the varied inheritance patterns, evidence types and phenotypic spectrum seen across hearing loss cohorts (see Variant Pilot methods below). The final set of specified rules is shown in **Table 1**.

**Table 1:** ACMG Criteria with HL-EP Specifications.

| **Pathogenic Criteria** |  |
| --- | --- |
| **Rule** | **Rule Description** |
| **PVS1** | Null variant in a gene with established LOF as a disease mechanism; see PVS1_Strong, PVS1_Moderate, PVS1_Supporting for reduced evidence applications |
| PVS1_Strong | See PVS1 flow chart for PVS1_Strong variants in gene where LOF is a known mechanism of disease |
| PVS1_Moderate | See PVS1 flowchart for PVS1_Moderate variants in gene where LOF is a known mechanism of disease |
| PVS1_Supporting | See PVS1 flowchart for PVS1_Supporting variants in gene where LOF is a known mechanism of disease |
| **PS1** | Same amino acid change as an established pathogenic variant; OR splice variants at same nucleotide and with similar impact prediction as previously reported pathogenic variant |
| **PS2** | 2 points per tables 5a and 5b:<br>Examples: 1 proven *de novo* occurrence; OR 2 assumed *de novo* occurrences |
| PS2_VeryStrong | 4 points per tables 5a and 5b:<br>Examples: 2 proven *de novo* occurrences; OR 1 proven + 1 assumed *de novo* occurrences; OR<br>4 assumed *de novo* occurrences |
| PS2_Moderate | 1 point per tables 5a and 5b:<br>Examples: 1 proven *de novo* occurrence (phenotype consistent but not specific to gene); OR<br>1 assumed *de novo* occurrence; OR 2 assumed *de novo* occurrences (phenotype/gene not specific) |
| PS2_Supporting | 0.5 points per tables 5a and 5b:<br>Example: 1 assumed *de novo* occurrence (phenotype/gene not specific) |
| **PS3** | Knock-in mouse model demonstrates the phenotype |
| PS3_Moderate | Validated functional studies show a deleterious effect (predefined list) |
| PS3_Supporting | Functional studies with limited validation show a deleterious effect |
| **PS4** | Fisher Exact or Chi-Squared analysis shows statistical increase in cases over controls, OR<br>Autosomal dominant: $\geq 15$ probands with variant, and variant meets PM2 |
| PS4_Moderate | Autosomal dominant: $\geq 6$ probands with variant, and variant meets PM2 |
| PS4_Supporting | Autosomal dominant: $\geq 2$ probands with variant, and variant meets PM2 |
| **PM1** | Mutational hot spot or well-studied functional domain without benign variation (*KCNQ4* pore-forming region; Gly residues in Gly-X-Y motifs of *COL4A3/4/5*) |
| **PM2** | Absent/Rare in population databases (absent or $\leq 0.00007$ (0.007%) for autosomal recessive, $\leq 0.00002$ (0.002%) for autosomal dominant) |
| PM2_<br>Supporting | Low MAF in population databases ($< 0.0007$ (0.07%) for autosomal recessive, |
| **PM3** | 1 point awarded from tables 7a and 7b<br>Example: Detected in trans with a pathogenic variant (recessive) |
| PM3_<br>VeryStrong | 4 points awarded from tables 7a and 7b<br>Example: Detected in trans in $\geq 4$ probands with a pathogenic variant (recessive) |
| PM3_<br>Strong | 2 points awarded from tables 7a and 7b<br>Example: Detected in trans in 2 probands with a pathogenic variant (recessive) |
| PM3_<br>Supporting | 0.5 points awarded from tables 7a and 7b<br>Examples: Two variants that meet PM2_Supporting detected in trans; OR a homozygous variant meeting PM2_Supporting |
| **PM4** | Protein length change due to an in-frame deletion or insertion that are not located in repetitive regions |
| **PM5** | Missense change at same codon as another pathogenic missense variant |
| PM5_<br>Strong | Missense change at same codon as two different pathogenic missense variants |
| **PM6** | See PS2 above |
| **PP1** | Segregation in one affected relative for recessive and two affected relatives for dominant |
| PP1_<br>Strong | Segregation in three affected relatives for recessive and five affected relatives for dominant |
| PP1_<br>Moderate | Segregation in two affected relatives for recessive and 4 affected relatives for dominant |
| ~~**PP2**~~ | ~~Missense in a gene with low rate of benign missense variants and pathogenic missense variants are common~~ |
| **PP3** | REVEL score $\geq 0.7$, or predicted impact to splicing using MaxEntScan |
| **PP4** | Patient's phenotype highly specific for gene or fully sequenced gene set (see specifications in Table 7) |
| ~~**PP5**~~ | ~~Reputable source without shared data classified variant as pathogenic~~ |
| **Benign Criteria** |  |
| **BA1** | MAF of $\geq 0.005$ (0.5%) for autosomal recessive; MAF of $\geq 0.001$ (0.1%) for autosomal dominant |
| **BS1** | MAF of $\geq 0.003$ (0.3%) for autosomal recessive; MAF of $\geq 0.0002$ (0.02%) for autosomal dominant. Likely benign, provided there is no conflicting evidence. |
| **BS1_Supporting** | MAF of $\geq 0.0007$ (0.07%) for autosomal recessive. No BS1_Supporting criteria for autosomal dominant. |
| **BS2** | Observation of variant (biallelic with known pathogenic variant for recessive) in controls inconsistent with disease penetrance. |
| **BS3** | ~~See BS3_Supporting~~ |
| **BS3_Supporting** | Functional study shows no deleterious effect (predefined list) |
| **BS4** | Non-segregation with disease |
| **BP1** | ~~Missense variant in a gene where only truncating variants cause disease~~ |
| **BP2** | Observed in trans with a dominant variant/observed in cis with a pathogenic variant (use with caution) |
| **BP3** | In-frame indels in repeat region without known function |
| **BP4** | Computational evidence suggests no impact; REVEL score $\leq 0.15$ or no impact to splicing in MaxEntScan. |
| **BP5** | Variant in an autosomal dominant gene found in a patient with an alternate explanation |
| **BP6** | ~~Reputable source without shared data classified variant as benign~~ |
| **BP7** | Silent variant with no predicted impact to splicing |

Strikethrough indicates rule was removed or not applicable. Abbreviations: MAF = minor allele frequency; Indels = insertion/deletions.

### Pathogenic allele frequency for recessive hearing loss

The frequencies of the most common pathogenic variants in HL were determined using a cohort of 3673 patients tested at the LMM. The three most common sequence variants were c.35delG, p.Val37Ile, and p.Met34Thr in *GJB2,* while the fourth variant was c.2299delG in the *USH2A* gene (**Table 2**). Test-based cohorts for the *GJB2* (n=1375 probands) or *USH2A* (n=1887 probands) genes were used to calculate the disease population allele frequency of each one of those variants.

**Table 2:** Most common pathogenic variants in hearing loss.

| Variant | LMM (Allele Frequency; Allele Ratio) | MORL (Allele Frequency) |
| --- | --- | --- |
| *GJB2*, c.35delG | 7.2% (198/2750) | 4.77% |
| *GJB2*, c.109G>A<br>p.(Val37Ile) | 4.4% (120/2750) | 2.01% |
| *GJB2*, c.101T>C<br>p.(Met34Thr) | 2.1% (57/2750) | 1.91% |
| *USH2A*, c.2299delG | 1.2% (46/3774) | 0.65% |

LMM: Partners Laboratory for Molecular Medicine; MORL: Iowa Molecular Otolaryngology and Renal Research Laboratories. The higher frequency for each of the above grey shaded variants was used to specify the population frequency rules. RefSeq transcripts are NM\_004004.5 for *GJB2* and NM\_206933.2 for *USH2A*.

The allele frequencies were confirmed using another HL cohort (n=2066) tested at the Molecular Otolaryngology and Renal Research Laboratories (MORL; Iowa City, IA). Although their frequencies were slightly different, the rankings of the four most common pathogenic variants was recapitulated in this cohort (**Table 2**). The frequency differences can be attributed to the fact that the MORL cohort excluded patients with positive findings upon *GJB2* prescreening. Therefore, the higher allele frequency (from the LMM) for each variant was conservatively used in calculating BA1 and BS1 for recessive HL. Our calculations were made under assumption of Hardy-Weinberg equilibrium and can be obtained using a recently published web application (https://www.cardiodb.org/allelefrequencyapp/)(Whiffin et al., 2017).

### Performance analysis of functional studies

To assess the performance characteristics (sensitivity, specificity, positive predictive value, negative predictive value) of the functional assays in each of *GJB2*, *SLC26A4*, and *COCH*, the number of true and false variant calls were determined as follows. Variants classified in ClinVar and included in functional assays were reviewed for classification accuracy and then used for informing performance. Pathogenic and likely pathogenic variants that had abnormal assay readout, defined as statistically significant deviation from wild type variant(s), were labeled as true positives. Normal assay readouts for pathogenic or likely pathogenic variants were considered false negatives. Similarly, benign and likely benign variants with normal assay readouts (i.e. does not significantly deviate from wild type variants) were counted as true negatives. False positives were benign or likely benign variants with abnormal assay readouts. A summary of true and false variant calls by five functional assays in the above three genes is shown in **Supplementary Table 1**.

### Variant Pilot

Following the specification of the ACMG/AMP guidelines, the rules were refined by interpreting a set of 51 variants in the *GJB2*, *SLC26A4*, *USH2A*, *MYO7A*, *CDH23*, *COCH*, *KCNQ4*, *MYO6,* and *TECTA* genes. Each variant was assessed independently by two variant curators, and the variant classification and rules applied were reviewed on a conference call to resolve discrepancies and reach consensus. Curators utilized ClinGen’s Variant Curation Interface (https://curation.clinicalgenome.org/) to assess and document the applicable rules for each variant.

## RESULTS

### Summary of Specifications

The HL-EP recommended specifications for 21 ACMG/AMP rules (**Table1**). Four rules had general recommendations on the application of the rule (PS1, PP3, BS4, BP4, and BP5). Seven rules had gene or disease based specifications (PS3, PM1, PM2, PP4, BA1, BS4, BP2). Seven rules had strength-level specifications (PVS1, PS2, PM3, PM5, PM6, PP1, BS3). Three rules had both gene/disease based specifications and strength-level specifications (PS4, BS1, BS2). No changes were recommended for three rules (PM4, BP3, and BP7), whereas four rules were considered not applicable (PP2, PP5, BP1, and BP6).

Two changes to the ACMG/AMP recommendations were made on how to combine criteria to classify sequence variants. The first is that a likely pathogenic classification can be reached if a variant in a gene associated with autosomal recessive HL meets PVS1 and PM2_Supporting. The second change was that a variant can be classified as likely benign if a variant meets BS1 without valid conflicting evidence that would suggest a pathogenic role.

It should be noted that we do not discuss the technical challenges associated with sequencing some of HL genes, such as homology, GC-rich and/or repetitive regions, which may affect variant calling and interpretation. False positive variants in these regions can exist in population or disease databases and may result in erroneous classifications. Laboratories should carefully assess the technical quality of any variant before its interpretation. A full list of technically challenging regions in 109 HL genes was recently characterized (DiStefano et al., 2018).

#### Population data (BA1, BS1, PM2)

Recessive and dominant forms of HL were distinguished with respect to the interpretation of variant frequency data from the general population. This is mainly due to differences in prevalence, penetrance, and gene contribution. Evidence-based estimates of these attributes are critical for establishing recessive and dominant allele frequency thresholds, at which a benign or pathogenic criterion of any strength level might be assigned for a given variant **Table 3**. For this work, the ExAC and gnomAD population databases were used, though other databases with a minimum of 2000 alleles are also sufficient. Caution should be used if the cutoffs specified below are applied on an uncharacterized population or a population with minimal representation in ExAC or gnomAD.

**Table 3:** Allele frequency cutoffs for dominant and recessive hearing loss disorders.

|  | ACMG-AMP Criteria | MAF | Preval. | Allelic Het.<sup>†</sup> | Penetrance |
| --- | --- | --- | --- | --- | --- |
| AUTOSOMAL<br>RECESSIVE | BA1 | ≥0.005<br>(0.5%) | 1/200 | 7.2% | 100% |
|  | BS1 | ≥0.003<br>(0.3%) | 1/200 | 4.4% | 100% |
|  | BS1_Supporting | ≥0.0007<br>(0.07%) | 1/200 | 1.0% | 100% |
|  | PM2_Supporting | <0.0007<br>(0.07%) | Can apply PM2_Supporting if MAF is < BS1_Supporting (0.07%) |  |  |
|  | PM2 | ≤0.00007<br>(0.007%) | Can apply PM2_moderate if MAF is an order of magnitude below BS1_Supporting (ie ≤0.007%); |  |  |
| AUTOSOMAL<br>DOMINANT | BA1 | ≥0.001<br>(0.1%) | 1/30 | 5% | 80% |
|  | BS1 | ≥0.0002<br>(0.02%) | 1/150 | 5% | 80% |
|  | PM2 | ≤0.00002<br>(0.002%) | Can apply PM2_Moderate if MAF is an order of magnitude below BS1 (ie ≤0.002%); |  |  |

Abbreviations: Preval. = Prevalence. Allelic Het. - Allelic heterogeneity. <sup>†</sup>Allelic heterogeneity encompassed both genetic and allelic heterogeneity because allele frequencies were derived from hearing loss cases spanning all genetic etiologies.

#### Recessive hearing loss

Based on a comprehensive literature search, a prevalence of 1 in 200 was used to specify the BA1 rule and to derive the BS1 and PM2 values for recessive HL. This was a highly conservative prevalence estimate from different ethnic groups and accounted for congenital and childhood onset HL (up to 19 years of age) with suspected genetic etiology (Lin, Niparko, & Ferrucci, 2011; Morton & Nance, 2006). Although HL prevalence increases with age, other contributing genetic (mostly dominant) and non-genetic (environmental, such as noise or drug related) factors may become more relevant in late-onset HL.

Penetrance was considered complete for recessive HL, based on the Expert Panel’s experience, with the exception of two variants, *GJB2* p.Met34Thr and p.Val37Ile, which were treated as exceptions. To derive conservative genetic and allelic heterogeneity values, the allele frequency of the most common pathogenic variants using HL patient cohorts tested at the LMM and MORL were calculated. The same four pathogenic variants were identified as most common among the two cohorts (See **Methods** and **Table 2**). The higher frequency value for each variant was used to derive a conservative minor allele frequency (MAF) threshold for BA1, BS1, and BS1_Supporting.

##### BA1 and BS1

Using an allele frequency web app, (https://www.cardiodb.org/allelefrequencyapp/), the MAF threshold for BA1 was set at ≥0.5% for autosomal recessive HL. This was calculated by using a prevalence of 1/200, complete penetrance (100%), and the allele frequency of the most common pathogenic variant found in cases (7.2% for c.35delG in *GJB2* - **Table 2**), Genetic heterogeneity was set to 1 given that the 7.2% allele frequency used was derived across all genetic causes of hearing loss. We then reviewed all reportedly pathogenic variants with frequencies near or higher than this threshold and developed a BA1 exclusion list which includes two low penetrance variants *GJB2* p.Met34Thr and p.Val37Ile as well as several founder mutations(**Supplementary Table 2**). Any variant exceeding 0.5% in any subpopulation, aside from variants on the exclusion list (**Supplementary Table 2**), would default to a benign classification. When implementing these allele frequency thresholds, a 95% confidence interval is used to determine the exact allele count for the cutoffs which allows accurate application to populations of varying size.

The MAF threshold for BS1 was set at ≥ 0.3% for recessive HL. This was calculated by using the allele frequency (4.4%) of the second most common pathogenic variant, p.Val37Ile in *GJB2* (**Table 2**), while maintaining the same prevalence, penetrance, genetic heterogeneity, and statistical correction used for the recessive BA1 derivation. Variants exceeding this frequency and having no other conflicting evidence for pathogenicity, meet the “Likely Benign” classification, provided that they are not the exclusion list in **Supplementary Table 2**.

The BA1 and BS1 cutoffs were validated using an independent approach, where an extensive curation of the literature was performed to estimate disease attributes (prevalence, gene contribution, pathogenic allele frequencies) of the most common causes of HL (*GJB2*, *SLC26A4*, *USH2A*, and *MYO7A*) across several ancestral groups globally (**Supplementary Methods** and **Supplementary Tables 3-6**). This approach validated the MAF thresholds determined above.

##### BS1_Supporting, PM2_Supporting and PM2

To enable broader use of population allele frequencies, we defined supporting strength levels of BS1 and PM2. BS1_Supporting was set at a MAF threshold of ≥0.07% using the same metrics as BA1 and BS1 (**Table 3**), but using a 1% allele frequency which is consistent with the most common pathogenic variant in the second most common recessive gene (c.2299delG in *USH2A* - **Table 2 - Table 2**) and above the frequency for most known pathogenic variants (Supplemental **Table 2**).

PM2 was set at ≤ 0.007%, which is an order of magnitude lower than the PM2_Supporting value (**Table 3**) and equivalent to an allele count of 14 in ExAC and 27 in gnomAD. Variants with frequencies between 0.07% and 0.007% can be awarded supporting evidence for pathogenicity (PM2_Supporting) (**Table 3**).

#### Dominant hearing loss

HL prevalence generally increases with age but with decreasing monogenetic contribution (Cunningham & Tucci, 2017). Most dominant HL forms are postlingual and therefore are more difficult to distinguish from genetic forms (Shearer, Hildebrand, & Smith, 1993). We chose prevalence in individuals aged 0-49 years to minimize those affected by age related HL. This age range is consistent with the age of onset of the major HL genes, including *TECTA* (Alloisio et al., 1999; Mustapha et al., 1999; Verhoeven et al., 1998), *DFNA5* (Bischoff et al., 2004; Booth, Azaiez et al., 2018; Van Laer et al., 1998), *COCH* (Nagy, Horvath, Trexler, Repassy, & Patthy, 2004; Robertson et al., 1998; Street et al., 2005), and *WFS1* (Eiberg et al., 2006; Fukuoka, Kanda, Ohta, & Usami, 2007; Hogewind et al., 2010). Based on an NHANES study (n=7490 patients), 1/15 individuals are expected to have HL by 49 years of age (Lin et al., 2011). For dominant BA1 derivation, we conservatively assumed that 50% of those individuals have a genetic cause, and calculated a prevalence of 1/30 (**Table 3**). This is a deliberate overestimation given that 50% is the estimated genetic contribution to congenital HL, which is the highest contribution among all age groups, and is therefore expected to be lower in adults.

Due to a lack of published data on dominant HL penetrance, we used an estimated penetrance of 80%, based on the collective research and clinical experience of its members. The relative contributions of known genes to dominant forms of HL were also assessed in a cohort of 2000 patients tested at MORL. Four genes, *TECTA*, *WFS1*, *KCNQ4*, and *COL11A2*, were the most common accounting for 20%, 13%, 13%, and 7%, respectively, of all dominant pathogenic variants. Furthermore, among all nonsyndromic dominant genes tested at the LMM (n=3076 probands), *KCNQ4* contributed the most for a total of four pathogenic and likely pathogenic variants in one proband each. Therefore, the maximum allelic heterogeneity for a dominant HL gene is conservatively 25%. Based on these data, we safely estimated that no dominant variant could contribute more than 5% of all HL in the 0-49 year age range. This is based on the assumption that the maximum contribution of one gene would be 20% (*TECTA*) and the maximum contribution of any one variant to a gene’s HL would be 25% (*KCNQ4*) (i.e. 20% × 25% = 5%). Our literature search also confirmed a maximum allelic heterogeneity of 5% in any given gene (Hildebrand et al., 2011; Iwasa, Nishio, & Usami, 2016; T. Naito et al., 2013).

Using the above conservative estimates, we set the dominant BA1 cutoff at ≥ 0.1% (**Table 3**). To derive the BS1 value, we used a less conservative estimate of dominant prevalence by assuming only 10% of all HL between ages 0-49 years is genetic. Therefore, a prevalence of 1/150 (1/15 × 10%) was used, while keeping all other values as in BA1, leading to a ≥ 0.02% BS1 cutoff. An allele frequency an order of magnitude less than this cutoff (i.e. ≤0.002%) was then considered sufficient to be awarded moderate evidence of pathogenicity (PM2). Using a 95% confidence interval, this is equivalent to 5 alleles in ExAC and 10 in gnomAD (**Table 3**).

It is important to note the phenotypic heterogeneity is exhibited by several of the HL genes. Genes known to cause both dominant and recessive forms of HL, such as *TECTA, COL11A2*, *GJB2* and *MYO7A* should be treated per the more conservative recessive rules for applying BA1 and BS1 rules since the frequency cutoffs for recessive inheritance are higher than those of dominant. This will help to avoid erroneously misclassifying variants as benign or likely benign. However, when considering pathogenicity evidence, the PM2 rule that matches the proposed form of HL (dominant vs recessive) should be applied.

X-linked forms of HL were not specifically addressed by the HL-EP. However, based on their relative rarity and the fact that most X-linked genes are not strictly recessive or dominant, with carrier females often displaying milder disease, the recessive thresholds for BA1, BS1, and BS1_Supporting can be applied for both X-linked recessive and X-linked dominant HL.

#### PS4, PS4_Moderate, PS4_Supporting: Prevalence in affected individuals statistically increased over controls

Published case-control studies demonstrating a statistically increased presence of a variant in affected individuals compared to race and ancestry-matched controls can be used as strong evidence towards pathogenicity, as outlined in the ACMG/AMP guidelines. For many variants, published case-control studies may not exist, but a case-control comparison to ancestry-matched controls in gnomAD can still provide critical evidence. This is particularly true for recessive variants with relatively high frequency in the general population. The following recommendations were made for variants in which a published case-control study does not exist.

A Chi-squared or Fisher’s Exact test using a 2×2 contingency table can be performed to determine if the MAF of a variant is statistically higher in cases compared to the general population. This analysis can be used as strong evidence if the MAF of the variant is significantly different (i.e. the p-value of the Chi-squared or Fisher’s Exact test is ≤0.05) in cases than in the general population. It is important to note that to do this comparison, the case and population alleles should be adequately ancestry-matched, the total numbers of positive and negative alleles should be known for both the case and the population cohorts, and the relevant gene must be definitively or strongly associated with hearing loss per the ClinGen Gene Curation framework (Strande et al., 2017).

Large general population databases, such as ExAC or gnomAD, are not true control cohorts and likely contain individuals with HL, given that HL is not screened out. As such, absence of statistical significance should not be used as evidence against pathogenicity.

The ACMG/AMP guideline allows for the counting of probands as evidence for PS4 in the absence of case-control studies. For autosomal recessive HL, this is achieved through PM3 (see below). For autosomal dominant HL, we adopted the specifications previously published by the ClinGen Cardiomyopathy Expert Panel (CMP-EP) given that dominant HL has similar attributes to cardiomyopathy in terms of penetrance and phenocopies (Kelly et al., 2018). These specifications state that if a variant is absent or extremely rare in the general population (i.e. PM2 is met), identification of ≥2, ≥6, or ≥15 probands with the same variant would reach supporting (PS4_Supporting), moderate (PS4_Moderate), or strong (PS4) evidence, respectively. The rule PM2 must be met to use these PS4 specifications. As noted above, PM2 for autosomal dominant HL was set at ≤0.002%, which is slightly lower than the PM2 value of ≤0.004% set by the CMP-EP.

## COMPUTATIONAL AND PREDICTIVE DATA

### Loss of function (LOF) variants (PVS1, PVS1_Strong, PVS1_Moderate, PVS1_Supporting)

The HL-EP and the ClinGen Sequence Variant Interpretation Working Group (SVI) group jointly refined this rule to be used for all recessive and haploinsufficient genes across most diseases including HL. This refinement included a detailed guideline for interpreting the PVS1 criteria across all LOF variant types (nonsense, frameshift, canonical splice site variants (+/-1,2), deletions, duplications, and initiation codon variants) while accounting for variant, exon and transcript-specific considerations and the varied evidence weights. This guideline is available in an accompanying paper in this issue (Abou Tayoun et al., this issue) and will not be discussed further here.

### Variants affecting the same amino acid residue (PS1, PM5, PM5_Strong)

The ACMG/AMP guideline considers the presence of a known pathogenic variant at the same amino acid residue as either strong or moderate evidence of pathogenicity. This PS1 rule is applied if the variant is a novel nucleotide change but results in the same amino acid change as the known pathogenic variant, and the PM5 rule is applied if the variant is a novel missense change at the same amino acid residue as a known pathogenic missense variant. It should be noted the ACMG/AMP guidelines recommend assessing whether the variants in question could have an impact at the DNA level, such as through splicing impacts, before applying PS1 or PM5. The HL-EP added a strength modification to the PM5 rule (PM5_Strong) so that a strong level of evidence can be applied if a novel missense variant is located at an amino acid residue with at least two other pathogenic variants at the same residue.

The HL-EP also specified that PS1 can be applied for DNA variants located at the same nucleotide in the splice consensus sequence as a known pathogenic variant. Based on the most conserved regions known to impact splicing, we defined the splice consensus sequence here as the first base and last three bases of the exon, the -12 through -1 positions in the intron, and the +1 through +6 positions in the intron. The +/-1,2 positions were excluded, given that these positions should be evaluated using PVS1 rules. In addition, splice prediction algorithms must predict a similar or greater effect than the comparator known pathogenic variant.

### Computational predictive tools (PP3, BP4, BP7)

Computational tools are commonly used to inform variant interpretation. For missense variants, the HL-EP recommends using REVEL, a computational tool that combines predictions from commonly used algorithms for missense variants (Ioannidis et al., 2016). REVEL was selected based on recently published work from members of ClinGen as well as evaluation within the HL-EP and the SVI (Ghosh, Oak, & Plon, 2017). Furthermore, we recommend specific REVEL scores are reached to apply supportive evidence of pathogenicity (PP3) or supportive evidence of benign (BP4). PP3 can be applied with a REVEL score of ≥0.7, given that 95% of benign variants are excluded at this score (Ioannidis et al., 2016). BP4 can be applied with a REVEL score of ≤0.15, given that 95% of pathogenic variants are excluded at this score (Ioannidis et al., 2016). Variants with a REVEL score between 0.15-0.7 will not receive either rule (PP3 or BP4).

The HL-EP recommends using MaxEntScan when assessing potential splicing impacts at the DNA level for missense, silent, and splice consensus sequence variants outside of the canonical ± 1 or 2 sites. For variants at the canonical splice sites, PP3 cannot be applied if PVS1 is used (Abou Tayoun *et al*, this issue).

### Computational and predictive data rules with no changes (PM4, BP3, BP7)

The HL-EP did not further specify the rules pertaining to protein length changing variants (PM4, BP3). However, the PVS1 flowchart mentioned above should be carefully followed for large in-frame deletions or insertions, particularly those affecting one or more exons. Silent variants with no predicted splicing impact can have BP7 applied, with no specified changes.

## FUNCTIONAL DATA

### Functional studies (PS3, PS3_Supporting, BS4_Supporting)

Based on systematic assessment of the literature, the HL-EP determined that no functional assays met a strong level of evidence for or against pathogenicity, except for a variant-specific knock-in mouse model. This is mainly due to the lack of well-established functional assays with enough data to assess their validity in predicting variant pathogenicity in most HL genes. Except for the functional assays in the *COCH*, *GJB2* and *SLC26A4* genes described below, a PS3_Supporting or BS3_Supporting can be appropriately used for evaluating variants in other genes not specified here if the assay is well-validated with positive and negative controls and the results are consistent with a known protein function.

#### GJB2

This gene, which encodes the gap junction protein, connexin 26, is the most common cause of genetic HL and has been well studied. Several research groups have measured the impact of genetic variants on its function primarily using electrical coupling or dye diffusion assays in *Xenopus* oocytes or mammalian cell systems (Bicego et al., 2006; Choung, Moon, & Park, 2002; D’Andrea et al., 2002; Lee, Derosa, & White, 2009; Mani et al., 2009). In both settings, a mutant and wildtype control cDNA are first transfected into separate oocytes or cell lines. In the electrical coupling assay, *GJB2* protein function is examined by altering the voltage in one cell and then measuring a current change in a neighboring cell through patch clamping or ion injection and dye sequestering (Ambrosi et al., 2013; Bruzzone et al., 2003; Haack et al., 2006; Mese, Londin, Mui, Brink, & White, 2004; Zhang et al., 2005). In the dye diffusion assay, movement of fluorescent dyes between neighboring cells is quantified as a measure of *GJB2* protein function. Variants disrupting *GJB2* function are expected to significantly reduce electrical coupling or dye transfer compared to control cDNA.

We searched the literature for variants that were assessed by these two assays, and that were validly classified as pathogenic, likely pathogenic, benign, or likely benign. We then determined the performance characteristics for each assay as described in the methods section and shown in **Supplementary Table 7**. Since both assays demonstrated a high sensitivity and positive predictive value, the HL-EP recommends using PS3_Moderate for any novel *GJB2* variant resulting in a statistically significant decrease or absence of current or dye transfer compared to wild-type run under the same experimental conditions. On the other hand, given the limited number of benign variants used as negative controls (**Supplementary Table 1**), the HL-EP was less confident regarding the ability of either assay to accurately predict benign effects, and in such cases, using BS3_Supporting is more appropriate.

It should be noted that in dye diffusion assays, a negative control (e.g. non-transfected or water injected control) should not display dye diffusion. In electrical coupling assays, water injected controls should display negligible currents at all potentials.

#### SLC26A4

The *SLC26A4* gene encodes an anion transporter, and its function has been assayed by measuring transport of radioactive anion isotopes (e.g. iodide or chloride) in cells expressing normal or mutant protein subunits (Bizhanova, Chew, Khuon, & Kopp, 2011; B. Y. Choi, Stewart et al., 2009; Dossena, Bizhanova et al., 2011; Dossena, Vezzoli et al., 2006; Gillam et al., 2004; Ishihara et al., 2010; Palos et al., 2008; Reimold et al., 2011; Scott et al., 2000; Yuan et al., 2012). Alternatively, other groups have used halide-sensitive fluorescent probes to measure anion transport (Cirello et al., 2012; Dossena, Bizhanova et al., 2011; Dossena, Nofziger et al., 2011; Dossena, Rodighiero et al., 2006; Fugazzola et al., 2007; Jang et al., 2014; Pera et al., 2008). An assay readout is considered abnormal if it is statistically different than the wild-type vector in the same experiment where a negative control (empty vector) shows absent immunofluorescent staining compared to the wild type. In both the radioactive and fluorescence based assays, several cell lines usually including *Xenopus* oocytes, mammalian, or human cells can be used to express the relevant *SLC26A4* gene variants. As with *GJB2*, we determined the performance characteristics of both assays by reviewing the results from valid pathogenic/likely pathogenic and benign/likely benign variants in ClinVar (**Supplementary Tables 1** and **7**). Again, there was a limited number of benign variants to assess the ability of both assays to predict benign effects (**Supplementary Table 1**). Furthermore, compared to *GJB2*, the two assays had lower positive predictive values (**Supplementary Table 7**). Therefore, we recommend using PS3_Supporting or BS3_Supporting if a pathogenic or a benign effect, respectively, was suggested by either assay.

#### COCH

This gene encodes for the Cochlin protein, which is normally secreted into the extracellular matrix of the inner ear (Fransen et al., 1999; Robertson et al., 2006; Robertson et al., 1998). Immunofluorescence and western blotting techniques have been used to assay cochlin secretion, localization, and post-translational modifications subsequent to expression in bacterial cells such as *E. coli* or mammalian cell lines (Bae et al., 2014; Cho et al., 2012; Grabski et al., 2003; Jung, Kim, Lee, Yang, & Choi, 2015). Compared to wild-type protein, abnormal patterns include absence of extracellular deposition (i.e. no secretion), intracellular aggregation (i.e. in the endoplasmic reticulum and Golgi), and stable dimerization due to inappropriate disulfide bond formation interfering with protein secretion and/or function (Cho et al., 2012; Nagy et al., 2004; Street et al., 2005; Yao, Py, Zhu, Bao, & Yuan, 2010). Based on the high sensitivity and positive predictive value (**Supplementary Tables 1 and 7**) of these protein assays, we recommend using PS3_Moderate if a variant exhibits an abnormal secretion, dimerization, or localization pattern compared to wild-type protein. On the other hand, due to a dearth of benign variants tested by these assays, we recommend using only BS3_Supporting if a variant shows a comparable pattern to wild type protein using these protein assays. No criteria should be applied if multiple assay results do not agree.

### Mutational hot-spots or functional domains (PM1)

The HL-EP identified one gene, *KCNQ4*, in the variant pilot set that harbored a functional domain for which PM1 can be applied. Missense variants located in the pore forming region (amino acids 270 - 297) of the *KCNQ4* gene are eligible for PM1, based on this region’s extreme intolerance to variation in the general population and significant enrichment of pathogenic missense variants observed in individuals with dominant sensorineural HL (T. Naito et al., 2013).

However, a systematic review of mutational hotspots or functional domains for all genes associated with hearing loss was not performed by the HL-EP. We acknowledge that PM1 can likely be applied to other such regions, such as variants impacting glycine residues in the Gly-X-Y motifs of the collagen genes (*COL11A2*, *COL4A3*, *COL4A4*, and *COL4A5*), which are essential for the triple helical structure formation (Chakchouk et al., 2015; I. Naito, Kawai, Nomura, Sado, & Osawa, 1996; Savige et al., 2016; Tryggvason, Zhou, Hostikka, & Shows, 1993).

## SEGREGATION DATA (PP1, PP1_Moderate, PP1_Strong, BS4)

Segregation with disease is used as evidence towards pathogenicity, and with an increasing number of segregations, stronger evidence can be applied. Given that segregation evidence is not specific to HL, the HL-EP worked in conjunction with the SVI to develop guidelines that further specify PP1. Three levels of evidence were recommended for both autosomal dominant and autosomal recessive segregations. Each strength level was based upon likelihood ratios of 4:1 (LOD 0.6), 16:1 (LOD 1.2), and 32:1 (LOD 1.5) to count as supporting, moderate, and strong evidence, respectively (**Table 4a**). For autosomal recessive segregations, unaffected individuals with equal probability of inheriting the variant(s) in question, typically siblings of a proband, can be taken into account, as shown in **Table 4b**.

**Table 4a:** Recommendations for PP1 (segregation evidence)

|  | General Recommendations |  |  |
| --- | --- | --- | --- |
|  | **<u>Supporting</u>** | **<u>Moderate</u>** | **<u>Strong</u>** |
| **Likelihood** | 4:1 | 16:1 | 32:1 |
| **LOD Score** | 0.6 | 1.2 | 1.5 |
| **Autosomal dominant threshold** | 2 affected segregations | 4 affected segregations | 5 affected segregations |
| **Autosomal recessive threshold** | See Table 4b | See Table 4b | See Table4b |

**Table 4b:** Recommendations for autosomal recessive segregation evidence (PP1)

|  |  | General Recommendations (Phenocopy not an issue) |  |  |  |  |  |  |  |  |  |  |
| --- | --- | --- | --- | --- | --- | --- | --- | --- | --- | --- | --- | --- |
|  |  | Unaffected Recessive Segregations |  |  |  |  |  |  |  |  |  |  |
|  |  | 0 | 1 | 2 | 3 | 4 | 5 | 6 | 7 | 8 | 9 | 10 |
| **Affected segregations** | **0** | 0 | 0.12 | 0.25 | 0.37 | 0.5 | 0.62 | 0.75 | 0.87 | 1 | 1.12 | 1.25 |
|  | **1** | 0.6 | 0.73 | 0.85 | 0.98 | 1.1 | 1.23 | 1.35 | 1.48 | 1.6 | 1.73 | 1.85 |
|  | **2** | 1.2 | 1.33 | 1.45 | 1.58 | 1.7 | 1.83 | 1.95 | 2.08 | 2.2 | 2.33 | 2.45 |
|  | **3** | 1.81 | 1.93 | 2.06 | 2.18 | 2.31 | 2.43 | 2.56 | 2.68 | 2.81 | 2.93 | 3.06 |
|  | **4** | 2.41 | 2.53 | 2.66 | 2.78 | 2.91 | 3.03 | 3.16 | 3.28 | 3.41 | 3.53 | 3.66 |
|  | **5** | 3.01 | 3.14 | 3.26 | 3.39 | 3.51 | 3.63 | 3.76 | 3.88 | 4.01 | 4.13 | 4.26 |
|  | **6** | 3.61 | 3.74 | 3.86 | 3.99 | 4.11 | 4.24 | 4.36 | 4.49 | 4.61 | 4.74 | 4.86 |
|  | **7** | 4.21 | 4.34 | 4.46 | 4.59 | 4.71 | 4.84 | 4.96 | 5.09 | 5.21 | 5.34 | 5.46 |
|  | **8** | 4.82 | 4.94 | 5.07 | 5.19 | 5.32 | 5.44 | 5.57 | 5.69 | 5.82 | 5.94 | 6.07 |
|  | **9** | 5.42 | 5.54 | 5.67 | 5.79 | 5.92 | 6.04 | 6.17 | 6.29 | 6.42 | 6.54 | 6.67 |
|  | **10** | 6.02 | 6.15 | 6.27 | 6.4 | 6.52 | 6.65 | 6.77 | 6.9 | 7.02 | 7.15 | 7.27 |

Affected segregations are counted in rows and unaffected segregations in columns. Affected segregations are affected family members in whom biallelic compound heterozygous or homozygous variants segregates. Unaffected segregations are defined as unaffected family members, typically siblings, who are at risk to inherit the two variants identified in the proband. These individuals should be either wild-type for both variants identified in the proband, or a heterozygous carrier for a single variant. Unaffected, carrier parents DO NOT count as unaffected segregations. There may be scenarios where individuals other than siblings could be counted as segregations, such as in families where one parent is affected with the autosomal recessive disorder, in large families with multiple branches, or in consanguineous families.
Each cell shows the LOD score of each combination of affected and unaffected segregations. LOD scores were calculated using a simplified LOD score formula, as described in Strande et al., 2017.

Non-segregations can be used as strong benign evidence (BS4), however this should be done with caution. For family members that have HL, but are negative for the variant(s) in question (phenotype-positive, genotype-negative), there is a possibility for a phenocopy. Indications of a potential phenocopy could include differences in phenotype, such as HL age of onset, severity, and the audiogram shape.

For family members that are positive for the variant(s) in question, but have normal hearing (genotype-positive, phenotype-negative), confounding factors that must be taken into consideration include age-related penetrance and variable expression. It is recommended that BS4 is only applied for genotype-positive, phenotype-negative scenarios for genes expected to have full penetrance, including most autosomal recessive genes. Furthermore, there must be confidence that the family member is truly unaffected. Diagnostic audiometric data, including ear specific audiograms or an auditory brainstem response (ABR) should be obtained. If there is any evidence for reduced penetrance, variable expression, or an age of onset later than the age of the unaffected person, BS4 should not be applied.

## DE NOVO OCCURRENCE (PS2, PS2_VeryStrong, PS2_Moderate, PS2_Supporting and PM6)

As defined by the 2015 ACMG guidelines, PM6 stands for "*de novo*, maternity and paternity not confirmed" and PS2 stands for "*de novo*, paternity and maternity confirmed". Approximately 80% of nonsyndromic HL is inherited in an autosomal recessive pattern with *de novo* occurrences being rare (Alford et al., 2014). However, *de novo* occurrences have been reported, particularly for genes associated with autosomal dominant or X-linked HL (Alvarez et al., 2003; J. W. Choi et al., 2015; Moteki et al., 2015; Yuan et al., 2009). Given that *de novo* strength-level specifications are not specific to HL genes, the HL-EP worked in conjunction with the SVI and adapted the current recommendations from SVI which are posted on the ClinGen website (https://www.clinicalgenome.org/working-groups/sequence-variant-interpretation/). Different levels of strength are applied to *de novo* occurrences depending on how many are observed, whether the phenotype is specific, and whether maternity and paternity are confirmed (**Table 5a and 5b**).

**Table 5a:** Points awarded per de novo occurrence(s)

| Phenotypic consistency | Points per Proband |  |
| --- | --- | --- |
|  | Confirmed de novo | Assumed de novo |
| Phenotype highly specific for gene | 2 | 1 |
| Phenotype consistent with gene but not highly specific | 1 | 0.5 |
| Phenotype consistent with gene but not highly specific and high genetic heterogeneity<sup>†</sup> | 0.5 | 0.25 |
| Phenotype not consistent with gene | 0 | 0 |

<sup>†</sup>Maximum allowable value of 1 may contribute to overall score

**Table 5b:** Recommendation for determining the appropriate ACMG/AMP evidence strength level for de novo occurrence(s)

| Supporting<br>(PS2_Supporting or PM6_Supporting) | Moderate<br>(PS2_Moderate or PM6) | Strong<br>(PS2 or PM6_Strong) | Very Strong<br>(PS2_VeryStrong or PM6_VeryStrong) |
| --- | --- | --- | --- |
| 0.5 | 1 | 2 | 1 |

## ALLELIC DATA (PM3, BS2)

### PM3, PM3_Strong, PM3_VeryStrong: In trans with a pathogenic variant (recessive disorders)

For recessive disorders, identifying a variant *in trans* with a pathogenic variant on the second allele is considered evidence towards pathogenicity. This was previously defined in the ACMG/AMP guidelines as moderate evidence (PM3) (Richards et al., 2015). However, if the variant under consideration is found in multiple probands with pathogenic variants on the other allele, the strength of that evidence should be considered greater. Therefore, in conjunction with the SVI, we developed a scoring system to account for the number of compound heterozygous and homozygous probands identified with a variant, as shown in **Table 6**. Each instance that a variant is found in a proband, either in homozygosity or compound heterozygosity with a second variant, is given a specified number of points, and the total points corresponds to the strength of PM3 that can be applied. Other scenarios, such as homozygous variants in consanguineous families, and variants of unknown phase, were also accounted for.

**Table 6a:** Default points for scoring variants that are observed in trans (PM3 rules)

| Classification/zygosity of other variant | Points per proband |  |
| --- | --- | --- |
|  | Known in trans | Phase unknown |
| Pathogenic/Likely pathogenic | 1.0 | 0.5 |
| Homozygous occurrence<br>(Max points from homozygotes=1.0) | 0.5 | N/A |
| Rare uncertain significance variant on other allele,<br>OR<br>Homozygous occurrence due to consanguinity,<br>(Max point= 0.5) | 0.25 | N/A |

**Table 6b:** Recommendation for determining the appropriate ACMG/AMP evidence strength level for in trans occurrence(s)

| PM3_Supporting | PM3<br>(Moderate) | PM3_Strong | PM3_Very Strong |
| --- | --- | --- | --- |
| 0.5 points | 1.0 points | 2.0 points | 4.0 points |

Furthermore, if a variant has been observed in many probands, each with a different pathogenic variant on the other allele, the likelihood that the variants are *in trans* is higher even if phasing has not been performed in every case, assuming the gene has been fully sequenced to detect all variants. In these cases, point values can be adjusted based on confidence of trans occurrence.

### BP2: Observation in trans with a pathogenic variant for dominant disorders or observation of a variant in cis with a pathogenic variant

The identification of a variant *in cis* with a pathogenic variant is supportive evidence for a benign interpretation. However, caution is recommended for the application of this rule for a case with an *in trans* observation of a dominant pathogenic variant. Careful assessment should include whether an earlier onset or more severe phenotype is possible, which could be consistent with an in trans pathogenic variant.

## PHENOTYPIC DATA (PP4 AND BP5)

The HL-EP developed recommendations for PP4, which, in the ACMG/AMP guideline, was intended as supporting evidence of pathogenicity when the patient’s phenotype is highly specific for a single gene. The HL-EP applied this rule to HL syndromes if all causative genes have been sequenced and the detection rate at least doubles when the added clinical feature is present. The rationale for using a doubling is based on a Bayesian model of the ACMG/AMP guideline whereby any supporting piece of evidence is equivalent to a doubling of likelihood (Tavtigian et al., 2018). The specific clinical features in addition to HL that are allowed for application of PP4 are shown in **Table 7**.

**Table 7:** Recommendations for specification of PP4.

| Gene | Syndrom<br>e | Phenotype | Detection rate in<br>unselected HL | Detection rate with<br>specified phenotype |
| --- | --- | --- | --- | --- |
| *SLC26A4* | Pendred<br>syndrome | Hearing loss with enlarged vestibular aqueduct (EVA) and/or Mondini malformation (incomplete partitioning type 2) | 2.6%<br>(Sloan-Heggen et al.,<br>2016) | 50% for a single<br>mutation<br>(Albert et al., 2006;<br>Azaiez et al., 2007;<br>Chattaraj et al., 2017; B.<br>Y. Choi, Madeo et al.,<br>2009; Pryor et al., 2005) |
| *MYO7A*,<br>*USH1C*,<br>*CDH23*,<br>*PCDH15*,<br>*USH1G* | Usher<br>syndrome<br>Type I | Moderately-severe to profound hearing loss and retinitis pigmentosa (onset typically in first decade), +/- vestibular dysfunction | 4.3%<br>(Sloan-Heggen et al.,<br>2016) | 78.7% for 2 mutations<br>(Le Quesne Stabej et al.,<br>2012) |
| *USH2A*,<br>*ADGRV1*<br>(GPR98),<br>*WHRN*<br>(DFNB31) | Usher<br>syndrome<br>Type II | Mild to severe hearing loss and retinitis pigmentosa (onset typically in first or second decade). | 2.9%<br>(Sloan-Heggen et al.,<br>2016) | 60.3%<br>(Le Quesne Stabej et al.,<br>2012) |
| *CLRN1* | Usher<br>syndrome<br>Type III | Progressive hearing loss with retinitis pigmentosa (variable onset) and vestibular dysfunction | <1%<br>(Sloan-Heggen et al.,<br>2016) | 50% (2/4) for 2<br>mutations<br>(Le Quesne Stabej et al.,<br>2012) |
| *WFS1* | LF –<br>SNHL | Low frequency autosomal dominant sensorineural hearing loss | 0.4% | 28.5% (2/7 cases, LMM<br>unpublished data) |
|  |  |  | (Sloan-Heggen et al., 2016) |  |
| *EYA1, SIX1* | Branchio-oto-renal syndrome | **<u>3 major – or – 2 major + 2 minor – or – 1 major + a 1st degree relative meeting criteria</u>**<br>**MAJOR:** Branchial anomalies, hearing loss/deafness, preauricular pits, renal anomalies<br>**MINOR:** External ear anomalies, middle ear anomalies, inner ear anomalies, preauricular tags, other (facial asymmetry, palate abnormalities) (Chang et al., 2004) | 0.2%<br>(Sloan-Heggen et al., 2016) | 45%<br>Testing must include deletion/duplication analysis<br>(Smith, 1993) |
| *OTOF, DFNB59* | ANSD | Auditory neuropathy spectrum disorder | 1%<br>(Sloan-Heggen et al., 2016) | 9-50%<br>(Matsunaga et al., 2012; Rodriguez-Ballesteros et al., 2008; Varga et al., 2006) |
| *MYH9* | MYH9-related disorders | Congenital macrothrombocytopenia and platelet macrocytosis (Pecci et al., 2014)<br>*NOTE: Patient must also be tested for DIAPH1 with no DIAPH1 variants identified* | <0.1%<br>(Sloan-Heggen et al., 2016) | >90%<br>(Pecci et al., 2014; Pecci et al., 2008; Savoia & Pecci, 1993) |
| *DIAPH1* | AD hearing loss with macrothrombocytopenia | Macrothrombocytopenia (Neuhaus et al., 2017; Stritt et al., 2016)<br>*NOTE: Patient must also be tested for MYH9 with no MYH9 variants identified*<br>*All truncating variants associated to AD occur in exon 27. Truncating variants in other exons have been associated with AR microcephaly.* | Detection rate is unknown.<br>Rare syndrome, prevalence <1/1000000<br>(DIAPH1-related sensorineural hearing loss-thrombocytopeniasyn<br>drome, n.d.) | Unknown |
| *PAX3*,<br>*SOX10*,<br>*MITF*,<br>*EDN3*,<br>*EDNRB* | Waardenburg syndrome | **<u>Two or more of the following:</u>**<br>1. Congenital SNHL<br>2. Pigmentary disturbances of iris<br>1. Complete heterochromia (two eyes of different color);<br>2. Partial or segmental heterochromia<br>3. Hypoplastic blue eyes – characteristic brilliant blue in both eyes<br>3. Pigmentary disturbances of hair<br>1. White forelock from birth or in teens<br>2. Premature graying before age 20<br>4. A first or second degree relative with 2 or more of criteria 1-3<br>5. *Dystopia canthorum* (*PAX3* variants)<br>6. *Hirschsprung disease* (*SOX10*, *EDN3*, *EDNRB*)<br>Modified from (Liu, Newton, & Read, 1995) | Detection rate is unknown.<br>Rare syndrome, prevalence of 9/100000<br>(Pingault et al. 2010; Pingault, 2015) | >75% for WS1 and WS3<br>30% for WS2<br>50% for WS4<br>(Pingault et al., 2010)<br>Testing must include deletion/duplication analysis |
| *POU3F4* | X-linked recessive deafness | Hearing loss + Incomplete partitioning type III (IAC dilation, fistulous connection between basal turn of cochlea and the IAC, cochlear hypoplasia, stapes fixation) | Detection rate unknown.<br>Prevalence is unknown. Inner ear abnormalities are pathognomonic.<br>(X-linked mixed deafness with<br>perilymphatic gusher, n.d.). | ≥26.7%<br>Testing must include deletion/duplication analysis including 5' promotor region. |
| *GPSM2* | Chudley-McCullough syndrome | Congenital severe to profound hearing loss with brain abnormalities including frontal polymicrogyria, grey matter heterotopia, cerebellar dysplasia, ventriculomegaly with small frontal horns, agenesis of the corpus callosum, arachnoid cysts (Doherty et al., 2012). | Detection rate unknown.<br>Rare. Prevalence is 1 / 1 000 000<br>(Chudley-McCullough syndrome, n.d.) | >90%<br>(Diaz-Horta et al., 2012; Doherty et al., 2012) |
| *KCNE1, KCNQ1* | Jervell and Lange-Nielsen syndrome | Congenital sensorineural hearing loss (must be bilateral and severe to profound) and a reproducible prolonged QTc interval $\geq 480$ msec<br>*Notes:*<br>- GeneReviews states most patients reported to date have QTc $\geq 500$ msec<br>- Schwartz Score: $\geq 3.5$ points = high probability of LQTS; QTc $\geq 480$ msec (3 points) plus congenital deafness (0.5 points)<br>- Upper limit of normal QTc is 440 msec for males and 460 msec for post-pubertal females.<br>(Alders, Bikker, & Christiaans, 1993) | Detection rate unknown.<br>Rare. Prevalence is 1-9 / 1 000 000<br>(Celano et al. 2009) | $\geq 90\%$<br>(Tranebjaerg, Samson, & Green, 1993) |

Abbreviations: LF-SNHL: Low frequency sensorineural hearing loss. WS1 = Waardenburg syndrome, type 1; WS2 = Waardenburg syndrome, type 2; WS3 = Waardenburg syndrome, type 3; WS4 = Waardenburg syndrome, type 4.

The HL-EP further specified the BP5 rule, which the ACMG/AMP guidelines defined as supporting evidence of benign when a variant is identified in a case with an alternate cause. The HL-EP recommends that BP5 is not invoked for variants in autosomal recessive genes, given that affected individuals are frequently identified as carriers of a pathogenic recessive variants in genes unrelated to the cause of their HL. For variants in autosomal dominant genes, the rule can be applied; however, caution should be taken if the observation is seen with a more severe phenotype than typical given that the presentation may be due to an additive effect of multiple dominant mutations (e.g. George et al., 2016).

## REMOVED AND NOT APPLICABLE RULES

The HL-EP recommends that PP2 (missense variants in genes with low rate of benign missense variation) and BP1 (missense variants in genes where only LOF variants cause disease) are not used. These are not applicable to any known genes associated with HL. In addition, the HL-EP supports the SVI’s decision to remove the two rules pertaining to variant classifications from reputable sources without evidence available (PP5 and BP6) based on the published rationale (Biesecker & Harrison, 2018).

## COMBINING CRITERIA RULES

The HL-EP recommends following the rules for combining criteria as outlined in the ACMG/AMP guidelines, with the addition of two modifications.

First, the HL-EP specified that PVS1 and PM2_Supporting can be combined to reach a variant classification of likely pathogenic for variants in autosomal recessive genes. The ACMG/AMP criteria require combining at least one moderate or two supporting criteria with a very strong criteria to reach a likely pathogenic classification. This was decided given that the HL-EP adopted the stringent criteria for evaluating predicted LOF variants as described above under PVS1 such that we feel that rare variants meeting at least PM2_Supporting have at least a 90% chance of being pathogenic.

The second addition that the HL-EP added to the rules for combining criteria was that variants meeting BS1 with no conflicting evidence can be classified as Likely Benign. This is consistent with approaches taken by the RASopathy and Cardiomyopathy Expert Panels (Gelb et al., 2018; Kelly et al., 2018).

Finally, per the new PVS1 rule recommendation (Abou Tayoun et al., this issue), we recommend against combining PVS1 with PP3 for the canonical ±1 or 2 splice bases given that PP3 is essentially applying predictive assumptions already built into PVS1.

## VARIANT PILOT OF THE HEARING LOSS SPECIFIED RULES

A set of 51 variants in the *GJB2*, *MYO7A*, *USH2A*, *CDH23*, *MYO6*, *KCNQ4*, *COCH*, *TECTA*, and *SLC26A4* genes were selected to test and refine the above specifications set by the HL-EP. These variants were either present in ClinVar, present on the exclusion list for BA1 and BS1, or identified via literature search. All the major rule types supporting pathogenicity were tested, except for the rules specifying *de novo* occurrences which are extremely rare within HL (**Figure 2a**). Dual curation was performed, and then consensus was reached following discussion for all classifications and their supporting rules among the HL-EP. In addition, several rules, including PP1, PS4, PP3, BP4, and BP1_Supporting, were iteratively refined based on this variant pilot. A summary of the genes, phenotype, associated inheritance pattern, and gene-specific rules are shown in **Table 8**.

**Table 8:** Variant pilot genes and applicable gene-specific ACMG/AMP criteria.

| **Gene** | **Disease, Inheritance** | **PVS1 Applicable** | **PM1: Mutational hot spot or well-studied functional domain** | **Functional Assays** | **Phenotype (PP4) Applicable** |
| --- | --- | --- | --- | --- | --- |
| **CDH23** | Usher syndrome, AR | Yes | N/A | N/A | Yes |
| **COCH** | Nonsyndromic HL, AD | N/A | N/A | Localization, secretion, and dimerization studies performed using immunofluorescence and Western blotting techniques | N/A |
| **GJB2** | Nonsyndromic SNHL, AR | Yes | N/A | Electrical coupling assays, dye transfer assays | N/A |
| **KCNQ4** | Nonsyndromic SNHL, AD |  | amino acids 271-292 | N/A | N/A |
| **MYO6** | Nonsyndromic SNHL, AD | Yes | N/A | N/A | N/A |
| **MYO7A** | Usher syndrome, AR | Yes | N/A | N/A | Yes |
| **SLC26A4** | Pendred syndrome, AR |  | N/A | Radio isotope and fluorescence assays | Yes |
| **TECTA** | Nonsyndromic SNHL, AD | N/A | N/A | N/A | N/A |
|  | Nonsyndromic SNHL, AR | Yes | N/A | N/A | N/A |
| **USH2A** | Usher syndrome, AR | Yes | N/A | N/A | Yes |

Abbreviations: AR= autosomal recessive; AD = autosomal dominant; N/A = not applicable

**Figure 1.**
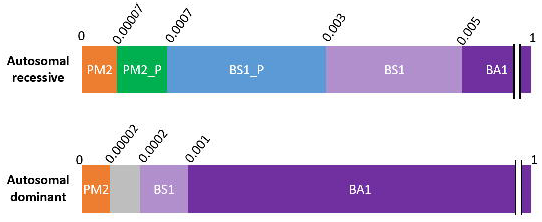
Population data criteria for autosomal recessive and autosomal dominant hearing loss. Minor allele frequencies are represented in decimals above each bar. Abbreviations:PM2_P = PM2_Supporting, BS1_P = BS1_Supporting.

**Figure 2.**
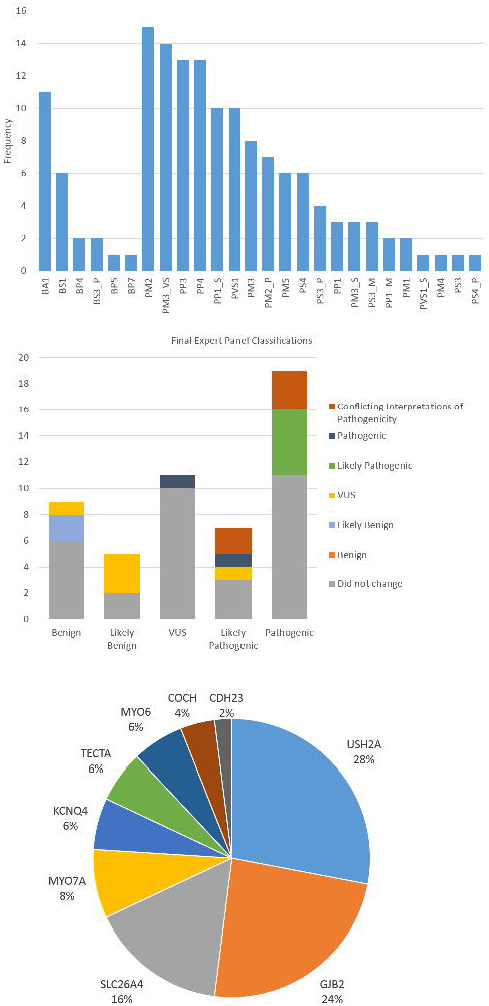
Classifications of 51 pilot variants. **A.** The frequency with which ACMG rules were applied during variant pilot. Rules applied with a modified strength are denoted by the rule followed by _P for Supporting, _M for Moderate, _S for Strong, and _VS for Very Strong. **B.** Final variant classifications of 51 pilot variants compared to initial classifications in ClinVar. The total height of each bar represents the total number of variants given each classification along the x-axis by the HL-EP. The colored segments of each bar represent variants whose ClinVar classification differs from the HL-EP classification. **C.** Distribution of pilot variants (n=51) by gene.

The list of the variants, their classifications and supporting rules are shown in **Supplementary Table 8**.

## DISCUSSION

Specifying the ACMG/AMP guidelines for HL posed unique challenges, which arose from high prevalence, high genetic heterogeneity, and multiple types of inheritance patterns. While published specifications exist for cardiomyopathy and RASopathy, these specifications focus on autosomal dominant disorders that typically have pathogenic missense variants as the causal variants. In contrast, HL is frequently autosomal recessive with LOF as the primary disease mechanism, and therefore, the HL-EP specifications addressed and utilized rules that have not been previously refined for more quantitative and nuanced use, including PM3 and PVS1. These rules were some of the most commonly used rules in the variant pilot and are likely to be frequent among all recessive disorders. As such, these specifications could be applied or serve as a groundwork for other autosomal recessive disorders that rely heavily on biallelic proband observations and predicted LOF variants.

Furthermore, a new combining rule for addressing novel LOF variants was created to address “under-classifying” variants that the HL-EP felt should be classified as likely pathogenic. By adding a rule whereby PM2_Supporting and PVS1 can be combined to reach a likely pathogenic variant classification, the HL-EP allowed convincing LOF variants with MAFs consistent with pathogenicity to be classified as likely pathogenic while remaining conservative for other potential rule combinations.

The HL-EP expects that these specifications will evolve as they are utilized by diagnostic laboratories, and as more information on the genetic basis of HL is identified, such as additional autosomal dominant loci, adult-onset HL, and founder mutations in under-studied cohorts. Furthermore, ongoing general refinements to the ACMG/AMP guidelines may be made by the ClinGen SVI which will need to be addressed.

The HL-EP is in the process of submitting for full Expert Panel status for its work on variant interpretation, with the goal of submitting expert variant classifications to ClinVar. These classifications will resolve discrepancies, move VUSs towards Benign or Pathogenic and increase the review status of hearing loss variant classifications present in ClinVar.

## Acknowledgments

*Authorship Contributions*

The work was overseen by the three co-chairs of the HL-EP: HLR, SSA and ANAT. AMO served as the coordinator for the HL-EP. The iterative ACMG/AMP rule specifications for discussion with the working group were developed and prepared by AMO, SEH, BJC, ARG, MTD, and RKS. AMO and ANAT created the first draft of the manuscript and SEH, ARG, MTD, RKS, LAS, KB, HA, KBA, MK, HLR, Yu Lu, Hatice Duzkale, Ignacio del Castillo, Minjie Luo, Xue Liu, Huijun Yuan, and Wenying Zhang provided critical edits and feedback. The final version was approved by all authors.

The following authors performed primary variant curations for the variant pilot: AMO, SEH, BJC, ARG, MTD, RKS, AC, NJB, LAS, JM, LH, KN, and JS. AG contributed to rules that impacted the SLC26A4 gene, including population based rules (BA1, BS1, PM2), functional evidence rules (PS3, BS3), and phenotype-specific rules (PP4). HK helped develop the population based rules (BA1, BS1, PM2) and the phenotype-specific rules (PP4). KB, HA, and KBA shared internal laboratory data that was instrumental in the development of the population data based rules and classification of pilot variants. MK and LAS provided their expert clinical knowledge.

The HL-EP would like to thank Steven Harrison, Tina Pesaran, Leslie G. Biesecker and the additional members from the SVI for their feedback on these specifications, and their contributions to the development of the PVS1, PM3, and PP1 rules.

The authors would also like to acknowledge additional members of the HLWG for their contributions, including Sonia Abdelhak, John Alexander, Zippora Brownstein, Rachel Burt., Byung Yoon Choi, Lilian Downie, Thomas Friedman, Anne Giersch, John Greinwald, Jeffrey Holt, Makoto Hosoya, Un-Kyung Kim, Ian Krantz, Suzanne Leal, Saber Masmoudi, Tatsuo Matsunaga, Matías Morín, Cynthia Morton, Hideki Mutai, Arti Pandya, Richard Smith, Mustafa Tekin, Shin-Ichi Usami, Guy Van Camp, Kazuki Yamazawa, Hui-Jun Yuan, Elizabeth Black-Zeigelbein, and Keijan Zhang.

